# Generative Modeling of Multi-mapping Reads with mHi-C Advances Analysis of High Throughput Genome-wide Conformation Capture Studies

**DOI:** 10.1101/301705

**Authors:** Ye Zheng, Ferhat Ay, Sündüz Keleş

## Abstract

Abstract Current Hi-C analysis approaches are unable to account for reads that align to multiple locations, and hence underestimate biological signal from repetitive regions of genomes. We developed and validated *mHi-C*, a *m*ulti-read mapping strategy to probabilistically allocate Hi-C multi-reads. mHi-C exhibited superior performance over utilizing only uni-reads and heuristic approaches aimed at rescuing multi-reads on benchmarks. Speciffically, mHi-C increased the sequencing depth by an average of 20% resulting in higher reproducibility of contact matrices and detected interactions across biological replicates. The impact of the multi-reads on the detection of signifficant interactions is influenced marginally by the relative contribution of multi-reads to the sequencing depth compared to uni-reads, cis-to-trans ratio of contacts, and the broad data quality as reflected by the proportion of mappable reads of datasets. Computational experiments highlighted that in Hi-C studies with short read lengths, mHi-C rescued multi-reads can emulate the effect of longer reads,. mHi-c also revealed biologically supported *bona fide* promoter-enhancer interactions and topologically associating domains involving repetitive genomic regions, thereby unlocking a previously masked portion of the genome for conformation capture studies.

## Introduction

DNA is highly compressed in the nucleus and organized into a complex three-dimensional structure. This compressed form brings distal functional elements into close spatial proximity of each other (***Dekker et al., 2002***; ***de Laat and Duboule, 2013***) and has far-reaching influence on gene regulation. Changes in DNA folding and chromatin structure remodeling may result in cell malfunction with devastating consequences (***Corradin et al., 2016***; ***Won et al., 2016***; ***Javierre et al., 2016***; ***Rosa-Garrido et al., 2017***; ***Spielmann et al., 2018***). Hi-C technique (***Lieberman-Aiden et al., 2009***; ***Rao et al., 2014***) emerged as a high throughput technology for interrogating the three-dimensional configuration of the genome and identifying regions that are in close spatial proximity in a genome-wide fashion. Thus, Hi-C data is powerful for discovering key information on the roles of the chromatin structure in the mechanisms of gene regulation.

There are a growing number of published and well-documented Hi-C analysis tools and pipelines (***Heinz et al., 2010***; ***Hwang et al., 2014***; ***Ay et al., 2014***a; ***Servant et al., 2015***; ***Mifsud et al., 2015***; ***Lun and Smyth, 2015***), and their operating characteristics were recently studied (***Ay and Noble, 2015***; ***Forcato et al., 2017***; ***Yardimci et al., 2017***) in detail. However, a key and common step in these approaches is the exclusive use of uniquely mapping reads. Limiting the usable reads to only uniquely mapping reads underestimates signal originating from repetitive regions of the genome which are shown to be critical for tissue specificity (***Xie et al., 2013***). Such reads from repetitive regions can be aligned to multiple positions (Figure 1A) and are referred to as multi-mapping reads or multi-reads for short. The critical drawbacks of discarding multi-reads have been recognized in other classes of genomic studies such as transcriptome sequencing (RNA-seq) (***Li and Dewey, 2011***), chromatin immunoprecipitation followed by high throughput sequencing (ChlP-seq) (***Chung et al., 2011***; ***Zenget al., 2015***), as well as genome-wide mapping of protein-RNA binding sites (CLIP-seq or RIP-seq) (***Zhang and Xing, 2017***). More recently, ***Sun et al. (2018)*** and ***Cournac et al. (2015)*** argued for a fundamental role of repeat elements in the 3D folding of genomes, highlighting the role of higher order chromatin architecture in repeat expansion disorders. However, the ambiguity of multi-reads alignment renders it a challenge to investigate the repetitive elements co-localization with the true 3D interaction architecture and signals. In this work, we developed mHi-C (Figure 1–Figure supplements 1 and 2), a hierarchical model that probabilistically allocates Hi-C multi-reads to their most likely genomic origins by utilizing specific characteristics of the paired-end reads of the Hi-C assay. mHi-C is implemented as a full analysis pipeline (https://github.com/keleslab/mHiC) that starts from unaligned read files and produces a set of statistically significant interactions at a given resolution. We evaluated mHi-C both by leveraging replicate structure of public of Hi-C datasets of different species and cell lines across six different studies, and also with computational trimming and data-driven simulation experiments.

**Figure 1.**
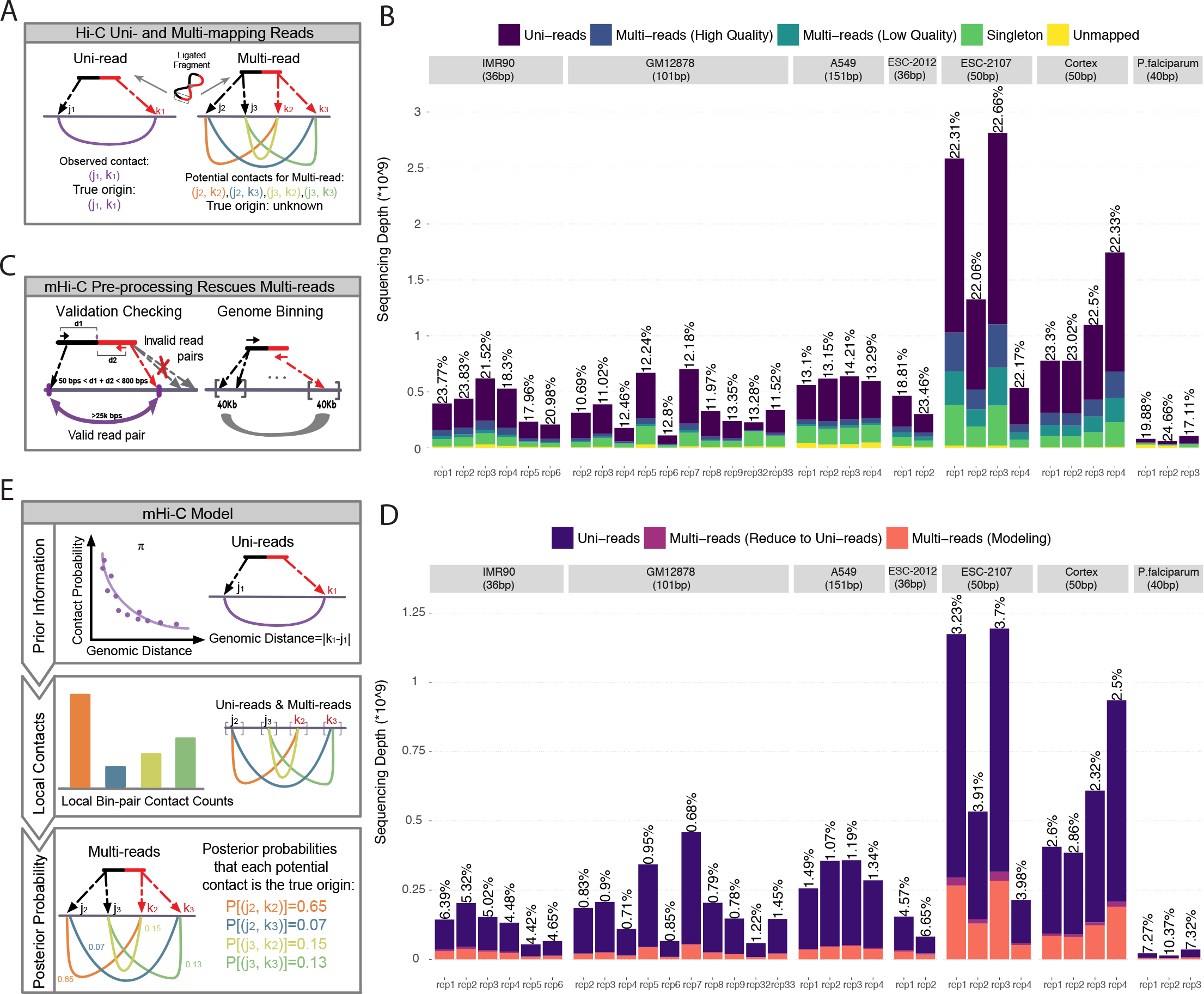
Overview of multi-reads and mHi-C pipeline. **(A)** Standard Hi-C pipelines utilize uni-reads while discarding multi-mapping reads which give rise to multiple potential contacts. **(B)** Total number of reads in different categories as a result of alignment to reference genome across the study datasets. Percentages of high-quality multi-reads compared to uni-reads are depicted on top of each bar. **(C)** Multi-mapping reads can be reduced to uni-reads within validation checking and genome binning pre-processing steps. **(D)** Aligned reads after validation checking and binning. Percentage improvements in sequencing depths due to multi-reads becoming uni-reads are depicted on top of each bar. **(E)** mHi-C modeling starts from the prior built by only uni-reads to quantify the relationship between random contact probabilities and the genomic distance between the contacts. This prior is updated by leveraging local bin pair contacts including both uni- and multi-reads and results in posterior probabilities that quantify the evidence for each potential contact to be the true genomic origin.

## Results

### Multi-reads significantly increase the sequencing depths of Hi-C data

For developing mHi-C and studying its operating characteristics, we utilized six published studies, resulting in eight datasets with multiple replicates, as summarized in Table 1 with more details in Figure 1–source data 1: Table 1. These datasets represent a variety of study designs from different organisms, i.e., human and mouse cell lines as examples of large genomes and three different stages of *Plasmodium falciparum* red blood cell cycle as an example of a small and AT-rich genome. Specifically, they span a wide range of sequencing depths (Figure 1 B), coverages and cis-to-trans ratios (Figure 1–Figure supplement 3), and have different proportions of mappable and valid reads (Figure 1–Figure supplement 4). Before applying mHi-C to these datasets and investigating biological implications, we first established the substantial contribution of multi-reads to the sequencing depth across these datasets with diverse characteristics. At read-end level (Appendix 1 Table 1 for terminology), after the initial step of aligning to the reference genome (Figure 1–Figure supplements 1 and 2), multi-reads constitute approximately 10% of all the mappable read ends (Figure 1–Figure supplement 5A). Moreover, the contribution of multi-reads to the set of aligned reads further increases by an additional 8% when chimeric reads (Appendix 1 Table 1) are also taken into account (Figure 1–source data 1: Table 2). Most notably, Figure 1–Figure supplement 6A demonstrates that, in datasets with shorter read lengths, multi-reads constitute a larger percentage of usable reads compared to uniquely mapping chimeric reads that are routinely rescued in Hi-C analysis pipelines (***Servant et al., 2015***; ***Lun and Smyth, 2015***; ***Durand et al., 2016***). Moreover, multi-reads also make up a significant proportion of the rescued chimeric reads (Figure 1–Figure supplement 6B). At the read pair level, after joining of both ends, multi-reads increase the sequencing depth by 18% to 23% for shorter read length datasets and 10% to 15% for longer read lengths, thereby providing a substantial increase to the depth before the read pairs are further processed into bin pairs (Figure. 1C; Figure 1–source data 1: Table 3).

**Table 1.**
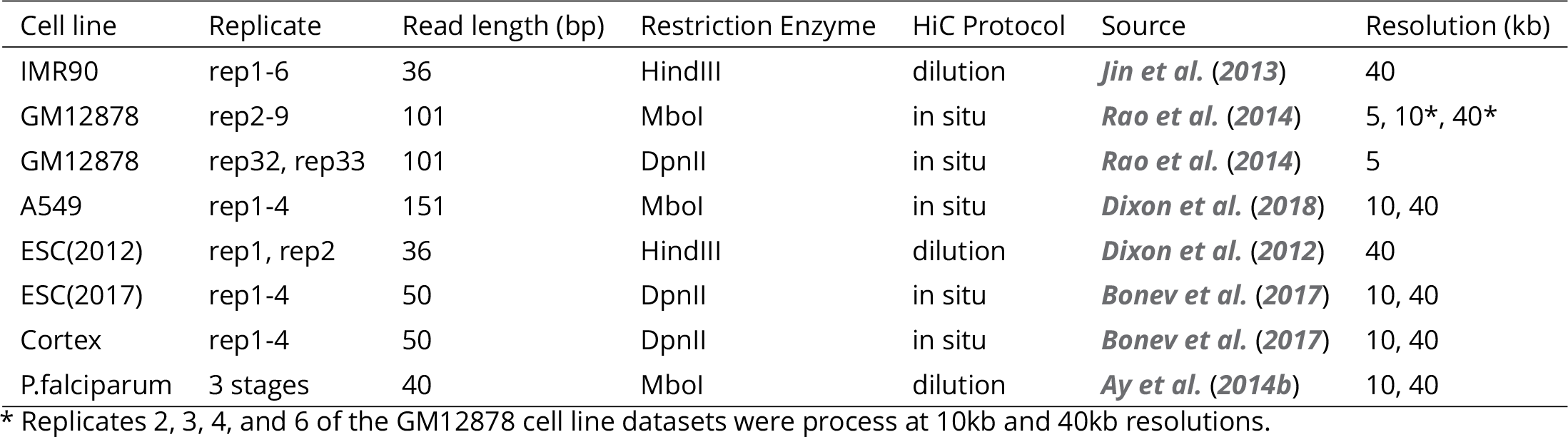
Hi-C Data Summary

### Multi-reads can be rescued at multiple processing stages of mHi-C pipeline

As part of the post-alignment pre-processing steps, Hi-C reads go through a series of validation checking to ensure that the reads that represent biologically meaningful fragments are retained and used for downstream analysis (Figure 1–Figure supplements 1 and 2, Appendix 1 Table 1). mHi-C pipeline tracks multi-reads through these processing steps. Remarkably, in the application of mHi-C to all six studies, a subset of the high-quality multi-reads are reduced to uni-reads either in the validation step when only one candidate contact passes the validation screening, or because all the alignments of a multi-read reside within the same bin (Figure 1C; Appendix 1 Table 1 and see Materials and Methods). Collectively, mHi-C can rescue as high as 6.7% more valid read pairs (Figure 1D) that originate from multi-reads and are mapped unambiguously without carrying out any multi-reads specific procedure for large genomes and 10.4% for *P. falciparum*. Such improvement corresponds to millions of reads for deeper sequenced datasets (Figure 1–source data 1: Table 4). For the remaining multi-reads (Figure 1D, colored in pink), which, on average, make up 18% of all the valid reads (Figure 1–Figure supplement 5B), mHi-C implements a novel multi-mapping model and probabilistically allocates them.

mHi-C generative model (Figure 1E and see Materials and Methods) is constructed at the bin-pair level to accommodate the typical signal sparsity of genomic interactions. The bins are either fixed-size non-overlapping genome intervals or a fixed number of restriction fragments derived from the Hi-C protocol. The resolutions at which seven cell lines are processed are summarized in Table 1. In the mHi-C generative model, we denote the observed alignment indicator vector for a given paired-end read *i* by vector ***Y_i_*** and use unobserved hidden variable vector ***Z_i_*** to indicate its true genomic origin. Contacts captured by Hi-C assay can arise as random contacts of nearby genomic positions or true biological interactions. mHi-C generative model acknowledges this feature by utilizing data-driven priors, ***π***_(*j,k*)_ for bin pairs *j* and *k*, as a function of contact distance between the two bins. mHi-C updates these prior probabilities for each candidate bin pair that a multi-read can be allocated to by leveraging local contact counts. As a result, for each multi-read i, it estimates posterior probabilities of genomic origin variable ***Z_i_***. Specifically, ***Pr***(***Z***_*i*,(*j,i*)_ = 1 | ***Y*_i_, π**) denotes the posterior probability, i.e., allocation probability, that the two read ends of multi-read ***i*** originate from bin pairs *j* and *k*. These posterior probabilities, which can also be viewed as fractional contacts of multi-read i, are then utilized to assign each multi-read to the most likely genomic origin. Our results in this paper only utilized reads with allocation probability greater than 0.5. This ensured the output of mHi-C to be compatible with the standard input of the downstream normalization and statistical significance estimation methods (***Imakaev et al., 2012***; ***Knight and Ruiz, 2013***; ***Ay et al., 2014***a).

### Probabilistic assignment of multi-reads results in more complete contact matrices and significantly improves reproducibility across replicates

Before quantifying mHi-C model performance, we provide a direct visual comparison of the contact matrices between Uni-setting and Uni&Multi-setting using raw and normalized contact counts. Figure 2A and Figure 2–Figure supplements 1-4 clearly illustrate how multi-mapping reads fill in the low mappable regions and lead to more complete matrices, corroborating that repetitive genomic regions are under-represented without multi-reads. In addition to increasing the sequencing depth in extremely low contact bins, higher bin-level coverage after leveraging multi-mapping read pairs appears as a global pattern across the genome (Figure 2–Figure supplement 5). For example, for the combined replicates of GM12878, 99.61% of the 5kb bins that potentially have interactions are covered by at least 100 contacts under the Uni&Multi-setting, compared to 98.72% under Uni-setting, thereby allowing us to study 25.55Mb more of the genome. These major improvements in coverage provide direct evidence that mHi-C is rescuing multi-reads that originate from biologically valid fragments.

**Figure 2.**
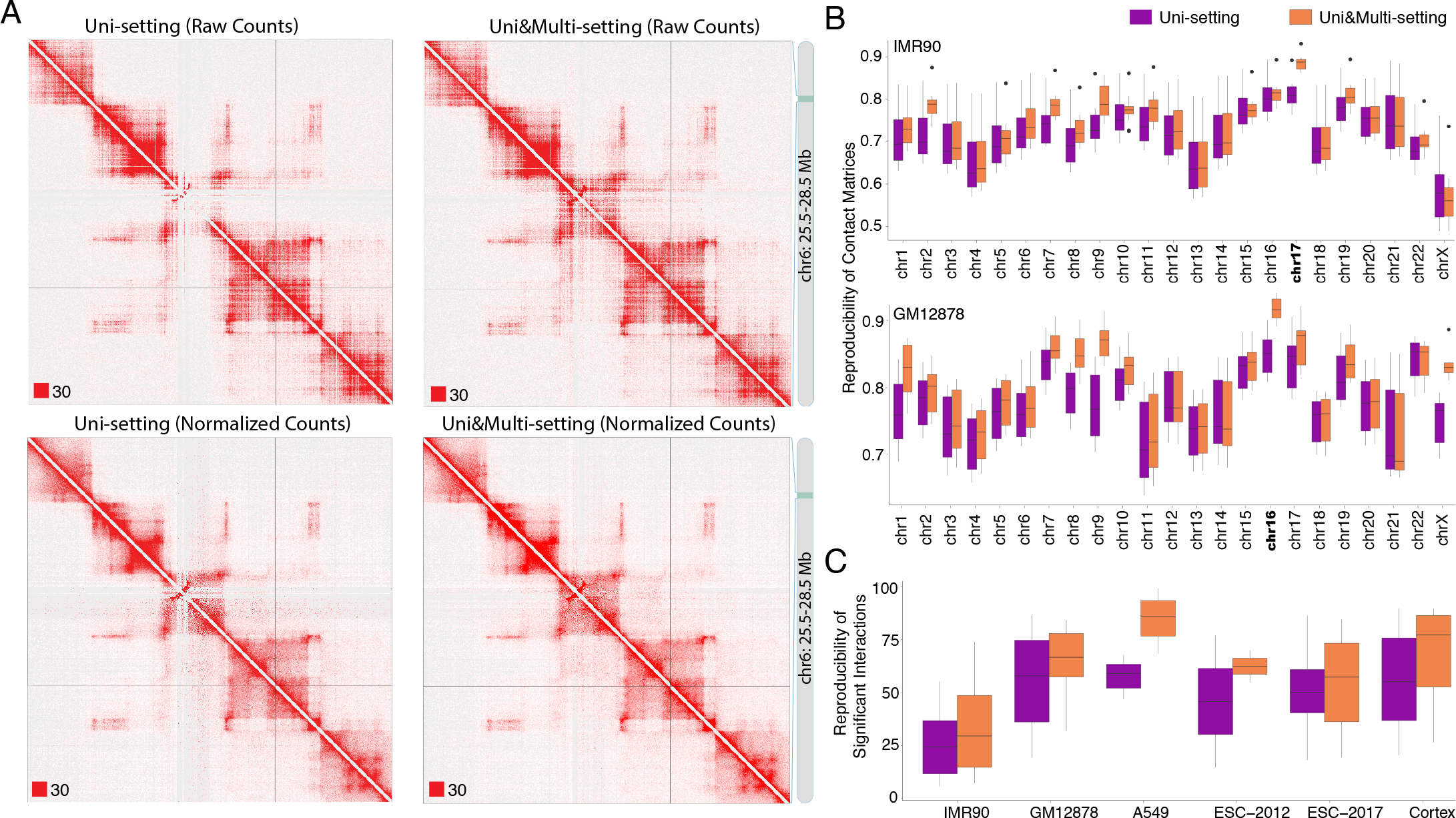
Global impact of multi-reads in Hi-C analysis. **(A)** Contact matrices of GM12878 with combined reads from replicates 2-9 are compared under Uni-setting and Uni&Multi-setting using raw and normalized contact counts for chr6:25.5Mb - 28.5Mb. White gaps of Uni-reads contact matrix, due to lack of reads from repetitive regions, are filled in by multi-reads, hence resulting in a more complete contact matrix. Such gaps remain in the Uni-setting even after normalization. Red squares at the left bottom of the matrices indicate the color scale. **(B)** Reproducibility of Hi-C contact matrices by HiCRep across all pairwise comparisons between replicates under the Uni- and Uni&Multi-settings (IMR90 and GM12878 are displayed). **(C)** Reproducibility of the significant interactions across replicates of the study datasets. Reproducibility is assessed by overlapping interactions detected at FDR of 5% for pairs of replicates within each study dataset.

We assessed the impact of multi-reads rescued by mHi-C on the reproducibility from the point of both raw contact counts and significant interactions detected. We used the stratum-adjusted correlation coefficient (***Yang et al., 2017***) for evaluating the reproducibility of Hi-C contact matrices. Figure 2B and Figure 2–Figure supplement 6 and 7 illustrate that integrating multi-reads leads to increased reproducibility and reduced variability of stratum-adjusted correlation coefficients among biological replicates across all the study datasets. Furthermore, we observe that, for some chromosomes, e.g., chr17 of IMR90 and chr16 of GM12878, the improvement in reproducibility stands out, without a systematic behavior across datasets. A close examination of improvement in reproducibility as a function of the ratio of rescued multi-reads to uni-reads across chromosomes highlights the larger proportion of multi-reads rescued for these chromosomes (Figure 2–Figure supplement 8).

In addition to the direct comparison of the raw contact matrices and their reproducibility, we identified the set of significant interactions by Fit-Hi-C (***Ay et al., 2014***a) and assessed the reproducibility of the identified interactions. Figure 2C shows that mHi-C significantly improves reproducibility of detected interactions across all the pairwise comparisons of replicates within each study dataset. Figure 2–Figure supplement 9A presents more details on the degree of overlap among the significant interactions identified at 5% and 10% false discovery rate (FDR) across replicates for the IMR90 datasets. These comparisons highlight that significant interactions specific to Uni&Multi-setting have consistently higher reproducibility than those specific to Uni-setting across all pairwise comparisons. Since random contacts tend to arise due to short genomic distances between loci, we stratified the significant interactions based on distance and reassessed the reproducibility as a function of the genomic distance between the contacts (Figure 2–Figure supplement 9B). Notably, significant interactions identified only by the Uni-setting and those common to both settings have a stronger gradual descending trend as a function of the genomic distance, indicating decaying reproducibility for long-range interactions. In contrast, Uni&Multi-setting maintains a relatively higher and stable reproducibility for longer genomic distances.

### Multi-reads detect novel signifcant interactions

At 5% false discovery rate, mHi-C detects 20% to 50% more novel significant interactions for relatively highly sequenced study datasets (Figure 3A and Figure 3–Figure source data 1; Figure 3A–Figure supplement 1 for other FDR thresholds and resolutions). The gains are markedly larger for datasets with smaller sequencing depths (e.g., ESC-2012) or extremely high coverage (e.g., *P. falciparum*). Overall gains in the number of novel contacts persist as the cut-off for mHi-C posterior probabilities of multi-read assignments varies (Figure 3–Figure supplement 2). At fixed FDR, significant interactions identified by the Uni&Multi-setting also include the majority of significant interactions inferred from the Uni-setting, indicating that incorporating multi-reads is extending the significant interaction list (low level of purple lines in Figure 3–Figure supplement 2).

**Figure 3.**
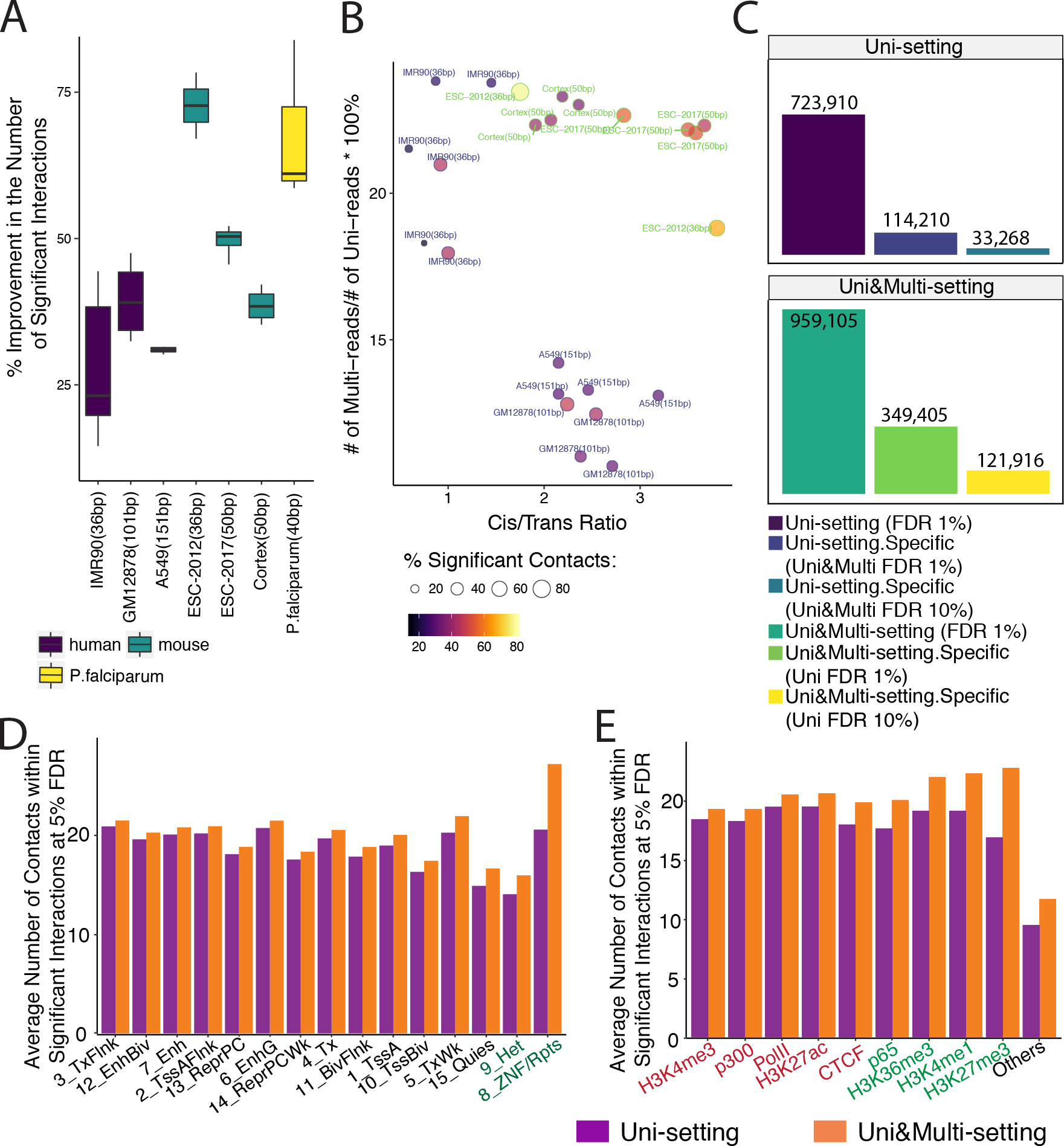
Gain in the numbers of novel significant interactions by mHi-C and their characterization by chromatin marks. **(A)** Percentage increase in detected significant interactions (FDR 5%) by comparing contacts identified in Uni&Multi-setting with those of Uni-setting across study datasets at 40kb resolution. **(B)** Percentage change in the numbers of significant interactions (FDR 5%) as a function of the percentage of mHi-C rescued multi-reads in comparison to uni-read and cis-to-trans ratios of individual datasets at 40kb resolution. **(C)** Recovery of significant interactions identified at 1% FDR by analysis at 10% FDR, aggregated over the replicates of GM12878 at 40kb resolution. Detailed descriptions of the groups are provided in Figure 3–Figure supplement 8. **(D)** Average number of contacts falling within the significant interactions (5% FDR) that overlapped with each chromHMM annotation category across six replicates of IMR90 identified by Uni- and Uni&Multi-settings. **(E)** Average number of contacts (5% FDR) that overlapped with significant interactions and different types of ChIP-seq peaks associated with different genomic functions (IMR90 six replicates). Red/Green labels denote smaller/larger differences between the two settings compared to the differences observed in the “Others” category that depict non-peak regions.

We leveraged the diverse characteristics of the study datasets and investigated the factors that impacted the gain in the detected significant interactions due to multi-reads. The top row of Figure 3–Figure supplement 3 summarizes the marginal correlations of the percentage change in the number of identified significant interactions (at 40Kb resolution and FDR of 0.05) with the data characteristics commonly used to indicate the quality of Hi-C datasets (excluding the high coverage *P. falciparum* dataset). These marginal associations highlight the significant impact of the relative contribution of multi-reads to the sequencing depth compared to uni-reads and cis-to-trans ratio of contacts (Figure 3–Figure supplement 4). Figure 3–Figure supplement 5 increase in the number of novel significant interactions for the GM12878 datasets in more detail across a set of FDR thresholds and at different resolutions, and includes two types of restriction enzymes. Specifically, Figure 3–Figure supplement 5C illustrates a clear negative association between the sequencing depth and the percent improvement in the number of identified significant interactions at 5kb resolution due to the larger impact of multi-reads on the smaller depth replicates. As an exception, we note that *P. falciparum* datasets tend to exhibit significantly higher gains in the number of identified contacts especially under stringent FDR thresholds (Figure 3A), possibly due to the ultra-high coverage of these datasets (Figure 3–Figure supplement 6). In addition to these marginal associations, Figure 3B and Figure 3–Figure supplement 7 display the percentage increase in the number of identified significant interactions as a function of the percentage increase in the real depth due to multi-reads and the cis-to-trans ratio across all the study datasets. A consistent pattern highlights that short read datasets with large proportion of mHi-C rescued multi-reads compared to uni-reads enjoy a larger increase in the number of identified significant interactions regardless of the FDR threshold, while for datasets with similar relative contribution of multi-reads, e.g., within lower depth IMR90, cis-to-trans ratios positively correlate with the increase in the number of identified significant interactions.

We next asked whether novel significant interactions due to rescued multi-reads could have been identified under the Uni-setting by employing a more liberal FDR threshold. Leveraging multi-reads with posterior probability larger than 0.5 and controlling the FDR at 1%, Fit-Hi-C identified 32.49% more significant interactions compared to Uni-setting (comparing dark green to dark purple bar in Figure 3C) and 36.43% of all significant interactions are unique to Uni&Multi-setting (light green bar over dark green bar in Figure 3C) collectively for all the four replicates of GM12878 at 40kb resolution. We observed that 34.89% of these novel interactions (yellow bar over the light green bar in Figure 3C) at 1% FDR (i.e., 12.71% compared to the all the significant interactions under Uni&Multi-setting) cannot be recovered even by a more liberal significant interaction list under Uni-setting at 10% FDR. Conversely, Uni&Multi-setting is unable to recover only 4.60% of the Uni-setting contacts once the FDR is controlled at 10% for the Uni&Multi-setting (light blue over dark purple bar in Figure 3C), highlighting again that Uni&Multi-setting predominantly adds on novel significant interactions while retaining contacts that are identifiable under the Uni-setting. A similar analysis for individual replicates of IMR90 are provided in Figure 3–Figure supplement 8 as well as those of GM12878 at the individual replicate level or collective analysis at 5kb, 10kb, and 40kb resolutions in Figure 3–Figure supplements 9-12. We further confirmed this consistent power gain by a Receiver Operating Characteristic (ROC) and a Precision-Recall (PR) analysis (Figure 3–Figure supplement 13). The PR curve illustrates that at the same false discovery rate (1-precision), mHi-C achieves consistently higher power (recall) than the Uni-setting in addition to better AUROC performance.

### Chromatin features of novel significant interactions

To further establish the biological implications of mHi-C rescued multi-reads, we investigated genomic features of novel contacts. Annotation of the significant interactions with ChromHMM segmentations from the Roadmap Epigenomics project (***Roadmap Epigenomics Consortium, 2015***) highlights marked enrichment of significant interactions in annotations involving repetitive DNA (Figure 3D, Figure 3–Figure supplement 14A). Most notably, ZNF genes & repeats and Heterochromatin states exhibit the largest discrepancy of the average significant interaction counts between the Uni- and Uni&Multi-settings. To complement the evaluation with ChromHMM annotations, we evaluated the Uni-setting and Uni&Multi-setting significant interaction enrichment of genomic regions harboring histone marks and other biochemical signals (The ENCODE Project Consortium, 2012; ***Jin et al., 2013***) (See Materials and Methods) by comparing their average contact counts to those without such signal (Figure 3E and Figure 3 – source data 2). Notably, while we observe that multi-reads boost the average number of contacts with biochemically active regions of the genome, they contribute more to regions that harbor H3K27me3 peaks (Figure 3E, Figure 3–Figure supplement 14B). Such regions are associated with downregulation of nearby genes through forming heterochromatin structure (***Ferrari et al., 2014***). Figure 3–Figure supplements 15-17 further provide specific examples of how increased marginal contact counts due to multi-reads are supported by signals of histone modifications, CTCF binding sites, and gene expression. Many genes of biological significance reside in these regions. For example, NBPF1 (Figure 3–Figure supplement 15) is implicated in many neurogenetic diseases and its family consists of dozens of recently duplicated genes primarily located in segmental duplications (***Safran et al., 2010***). In addition, RABL2A within the highlighted region of Figure 3–Figure supplement 16 is a member of RAS oncogene family.

### Multi-reads discover novel promoter-enhancer interactions

We found that a significant impact of multi-reads is on the detection of promoter-enhancer interactions. Overall, mHi-C identifies 14.89% more significant promoter-enhancer interactions at 5% FDR collectively for six replicates for IMR90 (Figure 4–source data 1: Table 1 and Figure 4 – source data 2). Of these interactions, 13,313 are reproducible among all six replicates under Uni&Multi-setting (Figure 4–source data 1: Table 2) and 62,971 are reproducible for at least two replicates (Figure 4–source data 1: Table 3) leading to 15.84% more novel promoter-enhancer interactions specific to Uni&Multi-setting. Figure 4A provides WashU epigenome browser (***Zhou et al., 2011***) display of such novel reproducible promoter-enhancer interactions on chromosome 1. Figure 4–Figure supplements 1-2 provides more such reproducible examples and Figure 4–Figure supplement 3 depicts the reproducibility of these interactions in more details across the 6 replicates.

**Figure 4.**
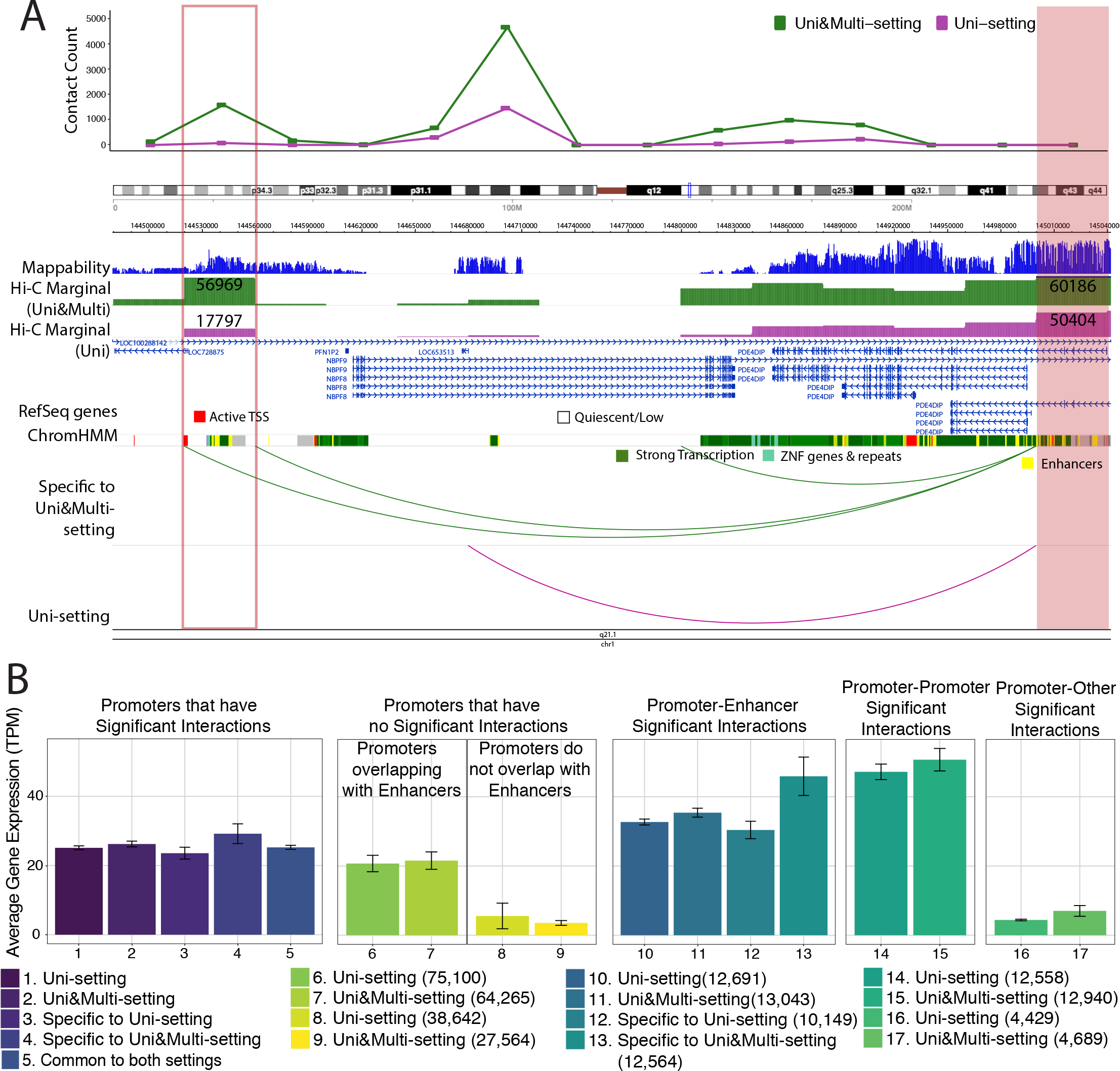
Novel promoter-enhancer interactions are reproducible and associated with actively expressed genes. **(A)** mHi-C identifies novel significant promoter-enhancer interactions (green arcs) that are reproducible among at least two replicates in addition to those reproducible under the Uni-setting (purple arcs). Shaded and the boxed regions correspond to the anchor and target bins, respectively. The top track displays the contact counts associated with the anchor bin under Uni- and Uni&Multi-settings. Related chromHMM annotation color labels are added around the track. The complete color labels are consistent with ChromHMM 15-state model at http://egg2.wustl.edu/roadmap/web_portal/chr_state_learning.html. **(B)** Average gene expression with standard errors for five different scenarios of interactions that group promoters into six different categories. In the first panel, significant interactions involving promoters are classified into five settings and the average gene expressions across genes with the corresponding promoters are depicted. The second panel involves two alignment settings and genes without any promoter interactions at 5% FDR. This panel is further separated into two categories: promoters that overlap with enhancer annotated regions and those that do not. The latter one serves as the baseline for average expression. Genes contributing to the third and fourth panel have promoter-enhancer, promoter-promoter interactions at 5% FDR. The fifth panel considers genes promoters of which have significant interactions with non-enhancer and non-promoter regions. Numbers in the parenthesis correspond to the number of transcripts in each category.

We next validated the novel promoter-enhancer interactions by investigating the expression levels of the genes contributing promoters to these interactions. Figure 4B supports that genes with significant interactions in their promoters generally exhibit higher expression levels (comparing bars 1-5 to bars 8-9 in Figure 4B). Furthermore, if these interactions involve an enhancer, the average gene expression can be 38.17% higher than that of the overall promoters with significant interactions (comparing bars 10-11 to bars 1-2 in Figure 4B). Most remarkably, newly detected significant promoter-enhancer interactions (bar 13 in Figure 4B) exhibit a stably higher gene expression level, highlighting that, without multi-reads, biologically supported promoter-enhancer interactions are underestimated. In addition, an overall evaluation of significant interactions (5% FDR) that considers interactions from promoters with low expression (TPM ≤ 1) as false positives illustrates that mHi-C specific significant promoter interactions have false positive rates comparable to or smaller than those of significant promoter interactions common to Uni- and Uni&Multi-settings (Figure 4–Figure supplement 4). In contrast, Uni-setting specific contacts have elevated false positive rates.

### Multi-reads refine the boundaries of topologically associating domains

We next investigated the impact of mHi-C rescued multi-reads on the topologically associating domains (TADs) (***Pombo and Dillon, 2015***), where we used a broad definition of TADs to include contact and loop domains. We used the DomainCaller (***Dixon et al., 2012***, ***2015***) to infer TADs of IMR90 datasets at 40kb resolution and Arrowhead (***Rao et al., 2014***) for GM12878 datasets at 5kb resolution under Uni&Multi-settings (Figure 5 – source data 1, 2). The detected TADs are compared to those under the Uni-setting. While this comparison did not reveal stark differences in the numbers of TADs identified under the two settings (Figure 5–Figure supplement 1), we found that Uni&Multi-setting identifies 2.01% more reproducible TADs with 2.36% lower non-reproducible TADs across replicates (Figure 5A). Several studies have revealed the role of CTCF in establishing the boundaries of genome architecture (***Ong and Corces, 2014***; ***Tang et al., 2015***; ***Hsu et al., 2017***). While this is an imperfect indicator of TAD boundaries, we observed that a slightly higher proportion of the detected TADs have CTCF peaks with convergent CTCF motif pairs at the boundaries once multi-reads are utilized (Figure 5–Figure supplements 2A-C). Figure 5B provides an explicit example of how the gap in the contact matrix due to depletion of multi-reads biases the inferred TAD structure. In addition to discovery of novel TADs (Figure 5–Figure supplement 3) by filling in the gaps in the contact matrix and boosting the domain signals, mHi-C also refines TAD boundaries (Figure 5–Figure supplements 4 and 5), and eliminates potential false positive TADs that are split by the contact depleted gaps in Uni-setting (Figure 5–Figure supplements 6-8). The novel, adjusted, and eliminated TADs are largely supported by CTCF signal identified using both uni- and multi-reads ChIP-seq datasets (***Zeng et al., 2015***) as well as convergent CTCF motifs (Figure 5–Figure supplement 2D), providing evidence for mHi-C driven modifications to these TADs and revealing a slightly lower false discovery rate for mHi-C compared to Uni-setting (Figure 5C, Figure 5–Figure supplement 2E, and Figure 5–Figure supplement 9).

**Figure 5.**
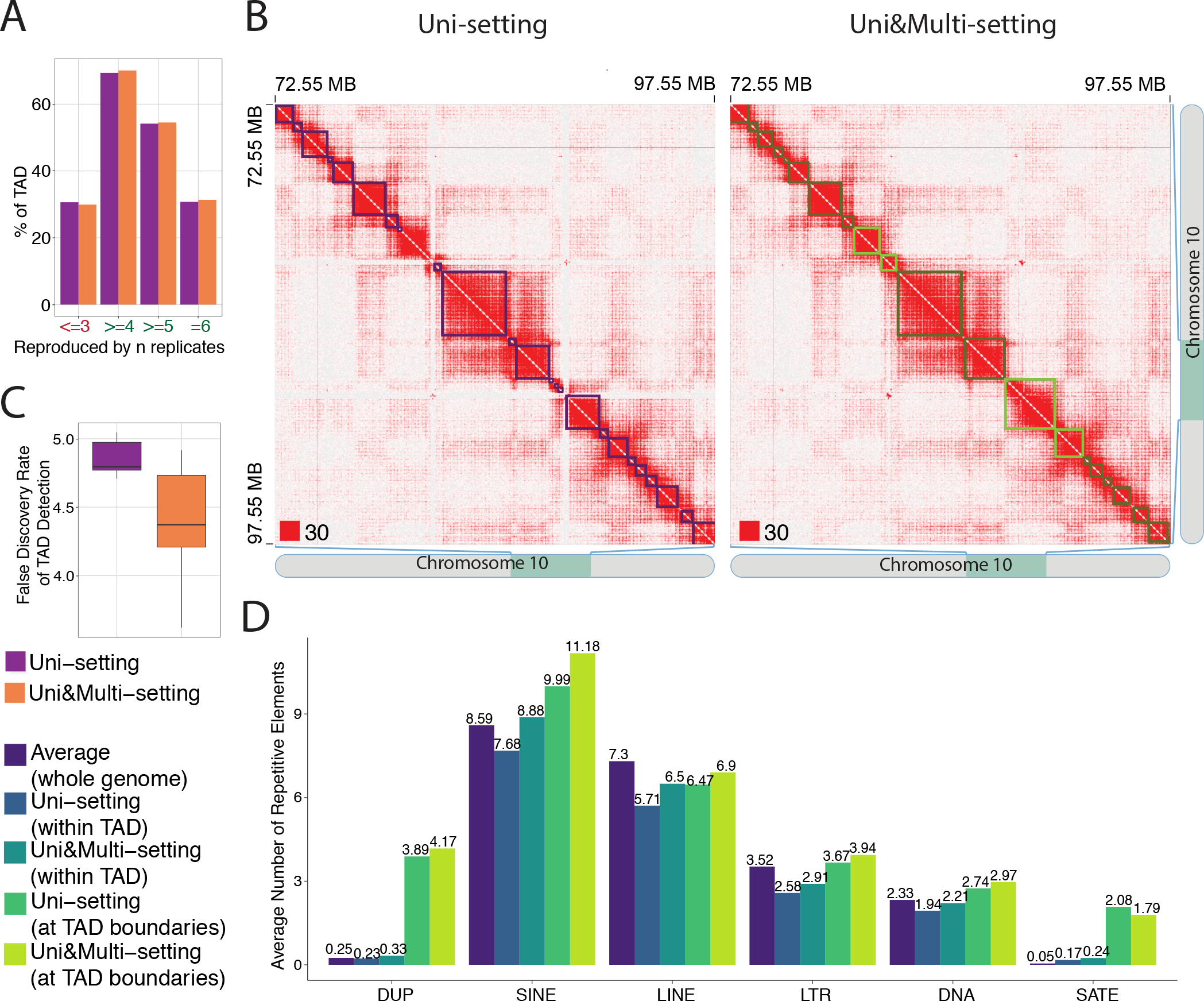
mHi-C rescued multi-reads refine detected topologically associating domains. **(A)** Percentage of topologically associating domains (TADs) that are reproducibly detected under Uni-setting and Uni&Multi-setting. TADs that are not detected in at least 4 of the 6 replicates are considered as non-reproducible. **(B)** Comparison of the contact matrices with superimposed TADs between Uni- and Uni&Multi-setting for chr10:72,550,000-97,550,000. Red squares at the left bottom of the matrices indicate the color scale. TADs affected by white gaps involving repetitive regions are highlighted in light green. Light green outlined areas correspond to new TAD boundaries. **(C)** False discovery rate of TADs detected under two settings. TADs that are not reproducible and lack CTCF peaks at the TAD boundaries are labeled as false positives. **(D)** Average number of repetitive elements at the boundaries of reproducible TADs compared to those within TADs and genomewide intervals of the same size for GM12878 at 5kb resolution.

Next, we assessed the abundance of different classes of repetitive elements, from the RepeatMasker (***Open, 2015***) and UCSC genome browser (***Casper et al., 2017***) hg19 assembly, at the reproducible TAD boundaries. Specifically, we considered segmental duplications (DUP), short interspersed nuclear elements (SINE), long interspersed nuclear elements (LINE), long terminal repeat elements (LTR), DNA transposon (DNA) and satellites (SATE). We utilized ±1 bin on either side of the edge coordinate of a given domain as its TAD boundary. At a lower resolution, i.e., 40kb for IMR90, each boundary is 120kb region and the percentages of TAD boundaries with each type of repetitive element illustrate negligible differences between the Uni-setting and Uni&Multi-setting (Figure 5–Figure supplement 10A). Similarly, due to the large sizes of the TAD boundaries, a majority of TAD boundaries harbor SINE, LINE, LTR, and DNA transposon elements. However, higher resolution analysis of the GM12878 dataset at 5kb reveals SINE elements as the leading category of elements that co-localizes with more than 99% of TAD boundaries followed by LINEs (Figure 5–Figure supplement 10B). This is consistent with the fact that SINE and LINE elements are relatively short and cover a larger portion of the human genome compared to other subfamilies (15% for SINE and 21% for LINE) (***Treangen and Salzberg, 2012***). We further quantified the enrichment of repetitive elements at TAD boundaries by comparing their average abundance with those within TADs and the genomic intervals of the same size across the whole genome as the baseline. Figure 5D and Figure 5–Figure supplement 11 show that SINE elements, satellites, and segmental duplications are markedly enriched at the TAD boundaries compared to the whole genome and within TADs. More interestingly, at higher resolution, i.e., 5kb for GM12878, the SINE category both have the highest average enrichment and is enhanced by mHi-C (Figure 5D). In summary, under uni&Multi-setting, the detected TAD boundaries tend to harbor more SINE elements supporting prior work that human genome folding is markedly associated with the SINE family ***Cournac et al. (2015)***.

### Large-scale evaluation of mHi-C with computational trimming experiments and simulations establishes its accuracy

Before further investigating the accuracy of mHi-C rescued multi-reads with computational experiments, we heuristic strategies for rescuing multi-reads at different stages of the Hi-C analysis pipeline as alternatives to mHi-C (Figure 6A; see Materials and Methods and Supplementary Text for the detailed description of the model-free approaches and related analysis). Specifically, AlignerSelect and DistanceSelect rescue multi-reads by simply choosing one of the alignments of a multi-read pair by default aligner strategy and based on distance, respectively. In addition to these, we designed a direct alternative to mHi-C, named SimpleSelect, as a model-free approach that imposes genomic distance priors in contrast to leveraging of the local interaction signals of the bins by mHi-C (e.g., local contact counts due to other read pairs in candidate bin pairs).

**Figure 6.**
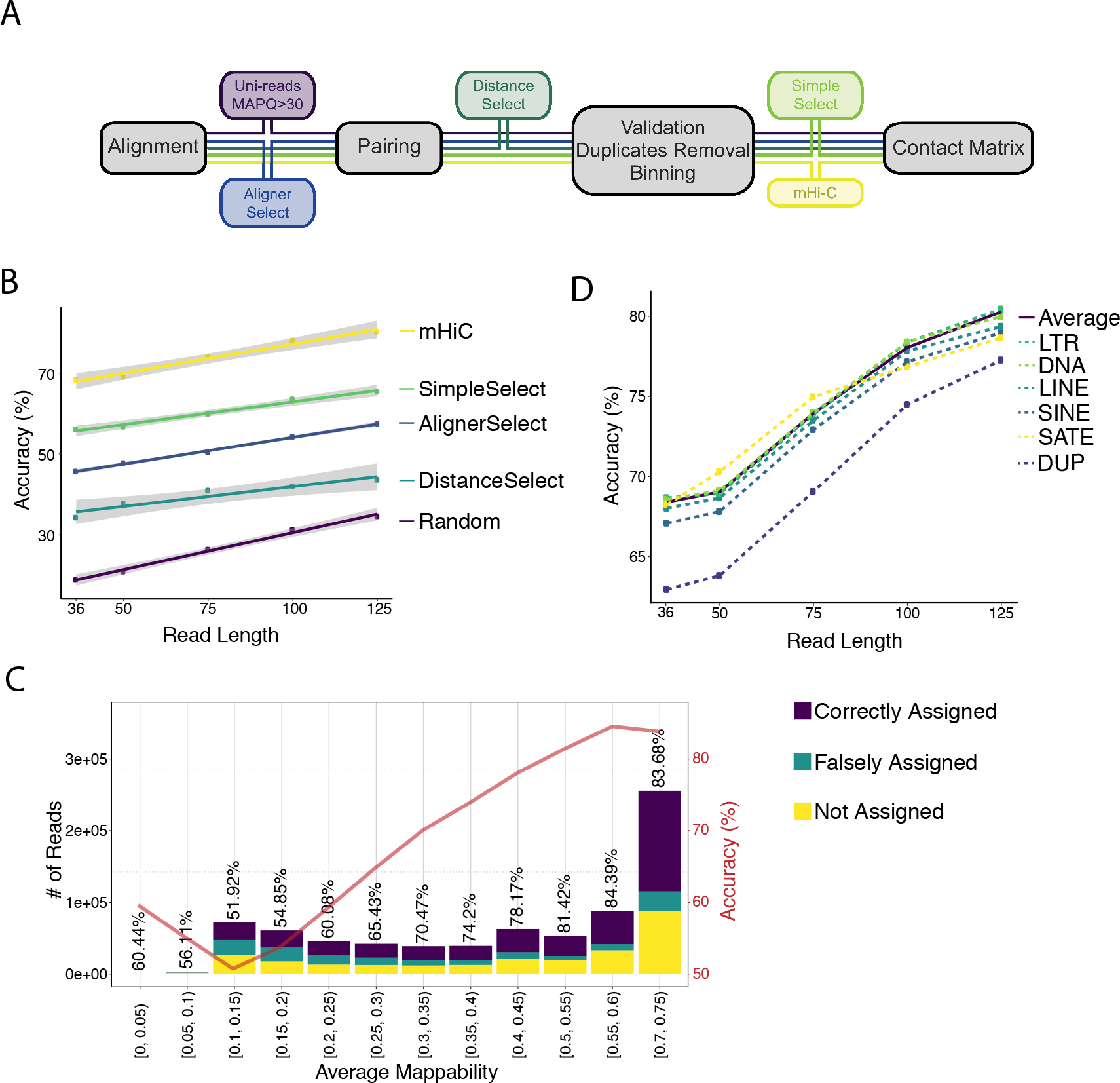
Assessing accuracy of mHi-C allocation by trimming experiments with the A549 study set of 151bp reads. **(A)** Intuitive heuristic strategies (AlignerSelect, DistanceSelect, SimpleSelect) for model-free assignment of multi-reads at various stages the of Hi-C analysis pipeline. **(B)** Accuracy of mHi-C in allocating trimmed multi-reads with respect to trimmed read length, compared with model-free approaches as well as random selection as a baseline. **(C)** Allocation accuracy with respect to mappability for 75bp reads. Red solid line depicts the overall accuracy trend. “Not assigned” category refers to multi-reads with a maximum posterior probability of assignment ≤ 0.5. **(D)** mHi-C accuracy among different repetitive element classes.

To evaluate the accuracy of mHi-C in a setting with ground truth, we carried out trimming experiments with the A549 151bp read length dataset and Hi-C data simulations where compared mHi-C to both a random allocation strategy as the baseline and the additional heuristic approaches we developed (Figure 6A). Specifically, we trimmed the set of 151bp uni-reads from A549 into read lengths of 36bp, 50bp, 75bp, 100bp, and 125bp. As a result, a subset of uni-reads at the full read length of 151bp with known alignment positions were reduced into multi-reads, generating gold-standard multi-read sets with known true origins. The resulting numbers of valid uni- and multi-reads are summarized in Figure 6–Figure supplement 1A in comparison with the numbers of valid uni-reads in the original A549 datasets. The corresponding multi-to-uni ratios of these settings vary with the lengths of the trimmed reads and their range covers the typical multi-to-uni ratios observed in the full read length datasets (Figure 6–Figure supplement 1B).

We first investigated the multi-read allocation accuracy with respect to trimmed read length, sequencing depth, and mappability at resolution 40kb. Figure 6B exhibits superior performance of mHi-C over both the model-free methods and the random baseline in correctly allocating multi-reads of different lengths to their true origins across intra- and inter-chromosomal contacts. As expected and illustrated by Figure 6B, the accuracy of multi-read assignment has an increasing trend with the read length. Specifically, it ranges between 70% and 90% for mHi-C and 20% and 35% for the random allocation strategy for the shortest and longest trimmed read lengths of 36bp and 125bp, respectively. When the allocated multi-reads are stratified as a function of the mappability, reads with the lowest mappability (<0.1) have accuracy levels of less than 32% to 70% across the trimmed read lengths (Figure 6C for 75bp, Figure 6–Figure supplement 2 for the other trimming lengths). Notably, the accuracy quickly reaches 74% to 87% for reads with mappability of at least 5 (Figure 6C, Figure 6–Figure supplement 2).

Next, we assessed the allocation accuracy among different classes of repetitive elements (Figure 6D). Allocations involving segmental duplication regions exhibit a systematically lower performance compared to other repeat classes and the average across the whole genome for all five trimming settings. Notably, even for these segmental duplication regions, the accuracy of mHi-C is markedly higher than both the model-free approaches and the random selection baseline displayed in Figure 6B. To finalize the accuracy investigation, we further varied the trimming setting by mixing uni-reads and multi-reads from different replicates (see setting (ii) of trimming strategies in Materials and Methods) and considering resolutions of 10kb and 40kb in addition to an empirical Hi-C simulation. Figure 6–Figure supplements 3-6 provide accuracy results closely following the results presented in this section from these additional settings and further validate significantly better performance of mHi-C compared to the random allocation and other heuristic approaches across different trimmed read lengths.

After establishing accuracy, we evaluated the impact of mHi-C rescued multi-reads of the trimmed datasets on the recovery of the (original) full read length contact matrices, topological domain structures, and significant interactions. To assess the recovery of the original contact matrix, we compared both the trimmed Uni- and Uni&Multi-settings with the gold standard Uni-setting at the full read length utilizing HiCRep (***Yang et al., 2017***). Figure 7A and Figure 7–Figure supplement 1 illustrate that mHi-C achieves significant improvement in the reproducibility across all chromosomes under all trimming settings compared to the Uni-setting. While the pattern of reproducibility with different read lengths in Figure 7A is consistent with the expectation that the longer trimmed reads yield contact matrices that are more similar to the full read length one, the improvement in reproducibility due to mHi-C is markedly larger compared to the gains from longer read sequences making multi-read rescue essential. For example, the reproducibility for Uni&Multi-setting at 50bp is 8.84% to 27.33% higher than that of Uni-setting at 125bp. TAD identification with these trimmed sets highlights the susceptibility of TAD boundary detection to the sequencing depth. Figure 7B displays an example where the trimmed Uni&Multi-setting achieved better recovery of TAD structure of the full-length dataset compared to trimmed Uni-setting. Overall, this performance is attributable to the accuracy of mHi-C assignments and the resulting increase in sequencing depth of the trimmed uni-read dataset. TAD analysis reveals that the improvement can be even more dramatic for short read contact matrices as illustrated in examples of Figure 7–Figure supplements 2-5. Finally, we compared the significant interactions detected by the trimmed Uni- and Uni&Multi-settings and observed that mHi-C rescued multi-reads in trimmed datasets enable detection of a larger number of interactions across a range of FDR thresholds (Figure 7–Figure supplement 6). Most notably, an evaluation of detection power for the top 10K significant interactions of the full-length dataset demonstrates that, while the Uni-setting can only recover 50% of these at the trimmed read length of 36bp, Uni&Multi-setting recovers 70% (Figure 7C and Figure 7–Figure supplement 7). We note that these power values are slightly underestimated because the full-length uni-read dataset also included chimeric reads that were rescued as uni-reads. In contrast, trimmed reads in the trimming experiments were generated from uni-reads without rescuing chimeric reads (see Materials and Methods; Figure 7–Figure supplement 8). Despite this, the 26.19% increase in sequencing depth due to multi-reads at the trimmed read length of 36bp (Figure 6–Figure supplement 1B) translated into a significantly better recovery of the significant interactions. Further assessment by ROC and PR analysis (Figure 7D, E and Figure 7–Figure supplement 9) of the set of significant contacts identified by both settings illustrates that Uni&Multi-setting exhibits these advantages without inflating the false discoveries. As reads get longer towards the full length, the ROC and PR curves converge under the two settings (Figure 7–Figure supplement 9).

**Figure 7.**
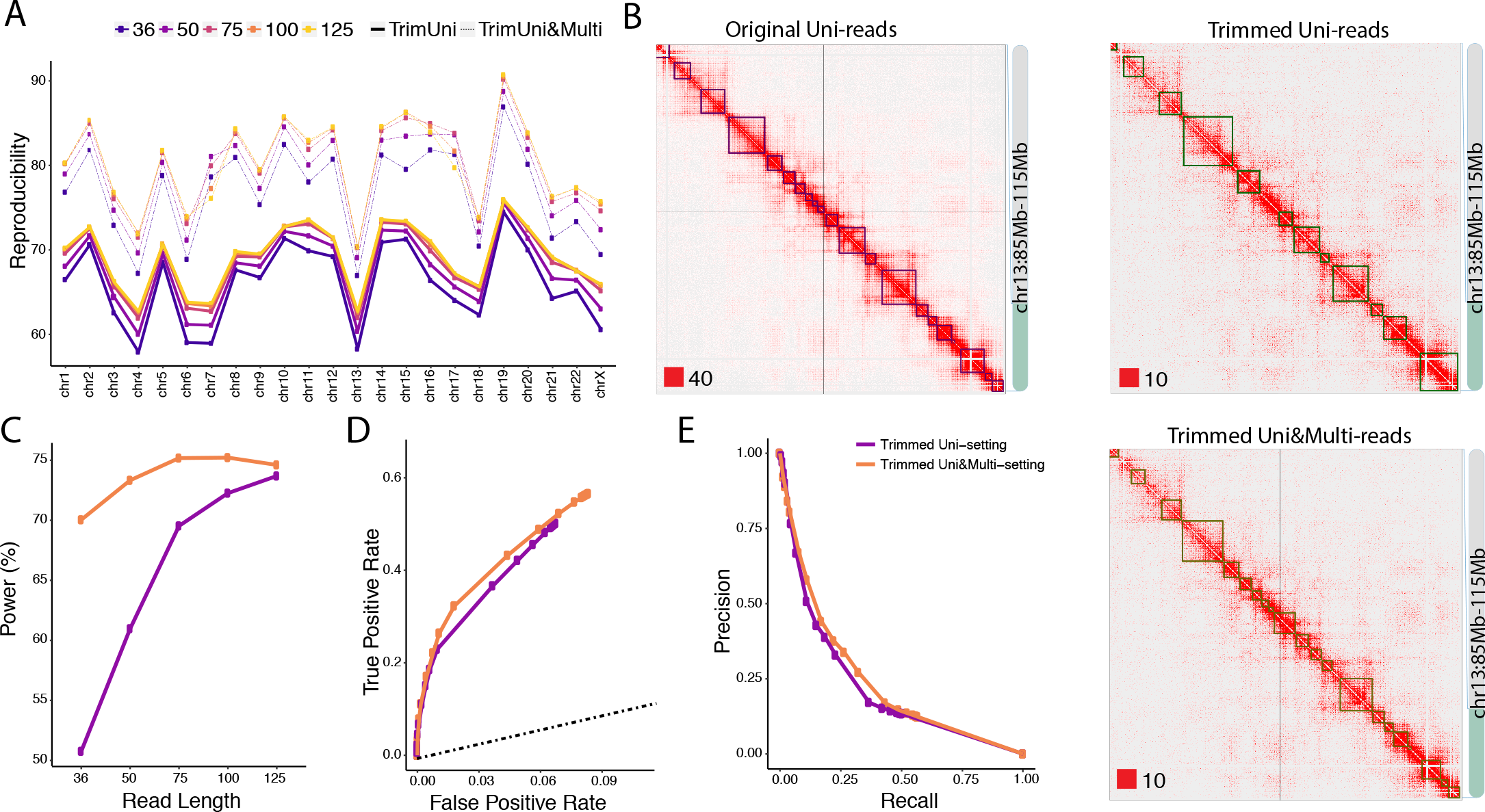
Trimmed uni- and multi-reads to recover the original contact matrix of the longer read dataset A549. **(A)** mHi-C rescued multi-reads of the trimmed dataset along with trimmed uni-reads lead to contact matrices that are significantly more similar to original contact matrices compared to only using trimmed uni-reads. **(B)** TAD detection on chromosome 13 with the original longer read uni-reads contact matrix, trimmed uni-reads (125bp) contact matrix, and trimmed uni- and multi-reads (125bp) contact matrix. **(C)** The power of recovering top 10,000 significant interactions of full read length dataset using trimmed reads under FDR10%. **(D, E)** Receiver Operating Characteristic (ROC) and Precision-Recall (PR) curves for trimmed Uni- and Uni&Multi-setting. The ground truth for these curves is based on the significant interactions identified by the full read length dataset at FDR of 10%. The dashed line is y = x.

## Discussion

Hi-C data are powerful for identifying long-range interacting loci, chromatin loops, topologically associating domains (TADs), and A/B compartments (***Lieberman-Aiden et al., 2009***; ***Yu and Ren, 2017***). Multi-mapping reads, however, are absent from the typical Hi-C analysis pipelines, resulting in under-representation and under-study of three-dimensional genome organization involving repetitive regions. Consequently, downstream analysis of Hi-C data relies on the incomplete Hi-C contact matrices which have frequent and, sometimes, severe interaction gaps spanning across the whole matrix. While centromeric regions contribute to such gaps, our results indicate that lack of multi-reads in the analysis is a significant contributor. Our Hi-C multi-read mapping strategy, mHi-C, probabilistically allocates high-quality multi-reads to their most likely positions (Figure 1E) and successfully fills in the missing chunks of contact matrices (Figure 2A and Figure 2–Figure supplements 1-4). As a result, incorporating multi-reads yields remarkable increase in sequencing depth which is translated into significant and consistent gains in reproducibility of the raw contact counts (Figure 2B) and detected interactions (Figure 2C). Analysis with mHi-C rescued reads identifies novel significant interactions (Figure 3), promoter-enhancer interactions (Figure 4), and refines domain structures (Figure 5). Our computational experiments with trimmed and simulated Hi-C reads validate mHi-C and elucidate the significant impact of multi-reads in all facets of the Hi-C data analysis. We demonstrate that even for the shortest read length of 36bp, mHi-C accuracy exceeds 74% (85% for longer trimmed reads) for regions with underlying mappability of at least 0.5 (Figure 6C and Figure 6–Figure supplement 2). mHiC significantly outperforms a baseline random allocation strategy as well as several other model-free and intuitive multi-read allocation strategies while achieving its worst allocation accuracy of 63% for reads originating from segmental duplications (Figure 6B and D). Trimming experiments further demonstrated the utility of multi-reads for contact matrix, TAD, and significant interaction recovery (Figure 7).

The default setting of mHi-C is intentionally conservative. In this default setting, mHi-C rescues high-quality multi-reads that can be allocated to a candidate alignment position with a high probability of at least 0.5. mHi-C allows relaxation of this strict filtering where instead of keeping reads with allocation probability greater than 0.5, these posterior allocation probabilities can be utilized as fractional contacts. We chose not to pursue this approach in this work as the current downstream analysis pipelines do not accommodate such fractional contacts. Currently, mHi-C model does not take into account potential copy number variations and genome arrangements across the genome. While mHi-C model can be extended to take into account estimated copy number and arrangement maps of the underlying sample genomes as we have done for other multi-read problems (***Zhang and Keleş, 2014***), our computational experiments with cancerous human alveolar epithelial cells A549 does not reveal any notable deterioration in mHi-C accuracy for these cells with copy number alternations.

## Materials and Methods

### mHi-C workflow

We developed a complete pipeline customized for incorporating high-quality multi-mapping reads into the Hi-C data analysis workflow. The overall pipeline, illustrated in Figure 1–Figure supplements 1 and 2, incorporates the essential steps of the Hi-C analysis pipelines. In what follows, we outline the major steps of the analysis to explicitly track multi-reads and describe how mHi-C utilizes them.

#### Read end alignment: uni- and multi-reads and chimeric reads

The first step in the mHi-C pipeline is the alignment of each read end separately to the reference genome. The default aligner in the mHi-C software is BWA (***Li and Durbin, 2010***); however mHi-C can work with any aligner that outputs multi-reads. The default alignment parameters are (i) edit distance maximum of 2 including mismatches and gap extension; and (ii) a maximum number of gap open of 1. mHi-C sets the maximum number of alternative hits saved in the XA tag to be to keep track of multi-reads. If the number of alternative alignments exceeds the threshold of 99 in the default setting, these alignments are not recorded in XA tag. We regarded these alignments as low-quality multi-mapping reads compared to those multi-mapping reads that have a relatively smaller number of alternative alignments. In summary, low-quality multi-mapping reads are discarded together with unmapped reads, only leaving uniquely mapping reads and high-quality multi-mapping reads for downstream analysis. mHi-C pipeline further restricts the maximum number of mismatches (maximum to be 2 compared to 3 in BWA default setting) to ensure that the alignment quality of multi-reads is comparable to that of standard Hi-C pipeline.

Chimeric reads, that span ligation junction of the Hi-C fragments (Figure 1–Figure supplement 1) are also a key component of Hi-C analysis pipelines. The ligation junction sequence can be derived from the restriction enzyme recognition sites and used to rescue chimeric reads. mHi-C adapts the pre-splitting strategy of diffHiC (***Lun and Smyth, 2015***), which is modified from the existing Cutadapt (***Martin, 2011***) software. Specifically, the read ends are trimmed to the center of the junction sequence. If the trimmed left 5′ ends are not too short, e.g., ≥ 25 bps, these chimeric reads are remapped to the reference genome. As the lengths of the chimeric reads become shorter, these reads tend to become multi-reads.

#### Valid fragment filtering

While each individual read end is aligned to reference genome separately, inferring interacting loci relies on alignment information of paired-ends. Therefore, read ends are paired after unmapped and singleton read pairs as well as low-quality multi-mapping ends (Figure 1–Figure supplement 1 and Appendix 1 Table 1) are discarded. After pairing, read end alignments are further evaluated for their representation of valid ligation fragments that originate from biologically meaningful long-range interactions (Figure 1–Figure supplement 1). First, reads that do not originate from around restriction enzyme digestion sites are eliminated since they primarily arise due to random breakage by sonication (***Belaghzal et al., 2017***). This is typically achieved by filtering the reads based on the total distance of two read end alignments to the restriction site. We required the total distance to be within 50-800 bps for the mammalian datasets and 50-500 bps for *P. falciparum*. The lower bound of 50 for this parameter is motivated by the chimeric reads with as short as 25 bps on both ends. Second, a single Hi-C interaction ought to involve two restriction fragments. Therefore, read ends falling within the same fragment, either due to dangling end or self-circle ligation, are filtered. Third, because the nature of chromatin folding leads to the abundance of random short-range interactions, interactions between two regions that are too close in the genomic distance are highly likely to be random interaction without regulatory implications. As a result, reads with ends aligning too close to each other are also filtered according to the twice the resolution rule. Notably, as a result of this valid fragment filtering, some multi-mapping reads can be counted as uniquely mapping reads (Appendix 1 Table 1 - 2b). This is because, although a read pair has multiple potential genomic origins dictated by its multiple alignments, only one of them ends up passing the validation screening. Once the multi-mapping uncertainty is eliminated, such read pairs are passed to the downstream analysis as uni-reads. We remark here that standard Hi-C analysis pipelines do not rescue these multi-reads.

#### Duplicate removal

To remove PCR duplicates, mHi-C considers the following two issues. First, due to allowing a maximum number of 2 mismatches in alignment, some multi-reads may have the exact same alignment position and strand direction with uni-reads. If such duplicates arise, uni-reads are granted higher priority and the overlapping multi-reads together with all their potential alignment positions are discarded completely. This ensures that the uni-reads that arise in standard Hi-C analysis pipelines will not be discarded as PCR duplicates in the mHi-C pipeline. Second, if a multi-mapping read alignment is duplicated with another multi-read, the read with smaller alphabetical read query name will be preserved. More often than not, if multi-read A overlaps multi-read B at a position, then it is highly likely that they will overlap at other positions as well. This convention ensures that it is always the read pair A alignments that are being retained (Figure 1–Figure supplement 2).

#### Genome binning

Due to the typically limited sequencing depths of Hi-C experiments, the reference genome is divided into small non-overlapping intervals, i.e., bins, to secure enough number of contact counts across units. The unit can be fix-sized genomic intervals or a fixed number of consecutive restriction fragments. mHi-C can switch between the two unit options with ease. After binning, the interaction unit reduces from alignment position pairs to bin pairs. Remarkably, multi-mapping reads, ends of which are located within the same bin pair, reduce to uni-reads as their potential multi-mapping alignment position pairs support the same bin pair contact. Therefore, there is no need to distinguish the candidate alignments within the same bin (Figure 1–Figure supplement 2 and Appendix 1 Table 1 - 3b).

#### mHi-C generative model and parameter estimation

mHi-C infers genomic origins of multi-reads at the bin pair level (Appendix 1 Table 1). We denoted the whole alignment vector for a given paired-end read *i* by vector ***Y_i_*** If the two read ends of read *i* align to only bin *j* and bin *k*, respectively, we set the respective components of the alignment vector as: *Y*_*i*(*j,k*)_ = 1 and *Y*_*i*, (*j*′, *k*′_ = 0, ∀ *j′* ≠ *j*,*k′* ≠ *k*. Index of read, *i*, ranges from 1 to *N*, where *N* is total number of valid Hi-C reads, including both uni-reads and multi-reads that pass the essential processing in Figure 1–Figure supplements 1 and 2. Overall, the reference genome is divided into *M* bins and *j* represents the bin index of the end, alignment position of which is upstream compared to the other read end position indicated by *k*. Namely, *j* takes on a value from 1 to *M* − 1 and *k* runs from *j* + 1 to the maximum value ***M***. For uniquely mapping reads, only one alignment is observed, i.e., 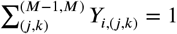. However, for multi-mapping reads, we have 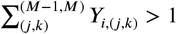.

We next defined a hidden variable *Z*_*i*,(*j,i*)_ to denote the true genomic origin of read *i*. If read *i* originates from position bin pairs *j* and *k*, we have *Z*_*i*,(*j,k*)_ = 1. In addition, a read can only originate from one alignment position pair on the genome; thus, 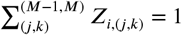 for both uni- and multi-reads. We define *O*_*i*_ = {(*j,k*): *Z*_*i*,(*j,k*)_ = 1} to represent true genomic origin of read *i* and 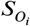 as the set of location pairs that read pair *i* can align to. Hence, ***Y***_*i*,(*j,k*)_ = 1, if 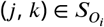. Under the assumption that the true alignment can only originate from those observed alignable positions, *O_i_* must be one of the location pairs in 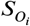. We further assume that the indicators of true origin for read *i*, ***Z_i_*** = (***Z***_*i*,(1,2)_,***Z***_*i*,(1,3)_, …, ***Z***_*i*,(*M*−1,*M*)_) are random draws from a Dirichlet - Multinomial distribution. Specifically,

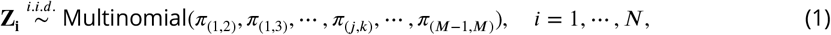

where *π*_(*j,k*)_ can be interpreted as contact probability between bin *j* and *k*(*j* < *k*). We further assume that

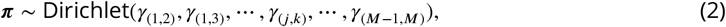

where ***π*** = (*π*_(1,2)_, *π*_(1,3)_, ···, *π*_(*M*−1,*M*)_) and *σ*_(*j,k*)_ is a function of genomic distance and quantifies random contact probability. Specifically, we adapt the univariate spline fitting approach from Fit-Hi-C (***Ay et al., 2014***a) for estimating random contact probabilities with respect to genomic distance and set *σ*_(*j,k*)_ = Spline(*j, k*) × *N* + 1. Here, *N* is the total number of valid reads as defined above and Spline(*j, k*) denotes the spline estimate of the random contact probability between bins *j* and *k*. Therefore, Spline(*j,k*)×*N* is the average random contact counts (i.e., pseudo-counts) between bin *j* and *k*. As a result, the probability density function of *π* can be written as:

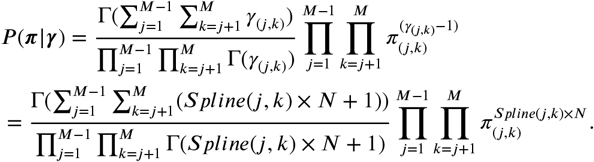

We next derive the full data joint distribution function.

##### Lemma 1

Given the true genomic origin under the mHi-C setting, the set of location pairs that read pair can align to will have observed alignments with probability 1.

**Proof**.

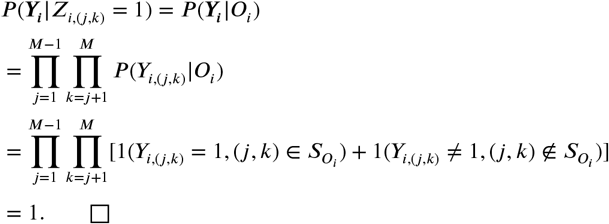

Based on Lemma 1, we can get the joint distribution ***P***(**Y**, **Z**|π) as

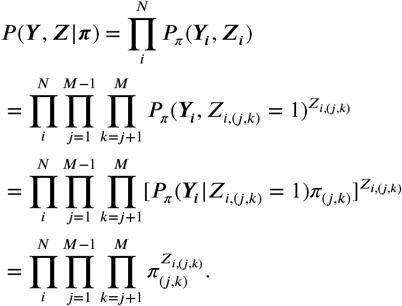

Using the Dirichlet-Multinomial conjugacy, we derive the posterior distribution of *π* as

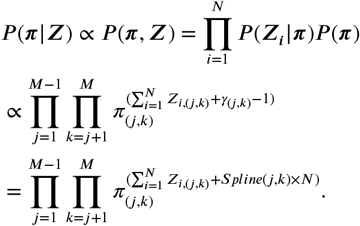

We next derive an Expectation-Maximization algorithm for fitting this model.

**E-step.**

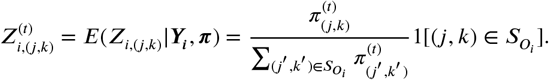

**M-step**.

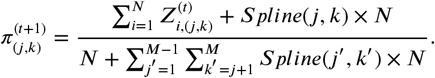

Estimate of the contact probability *π*_(*j,k*)_ in the M-step can be viewed as an integration of local interaction signal, encoded in 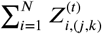 and random contact signal due to prior, i.e., Spline(*j, k*)×*N*.

The by-products of the EM algorithm are posterior probabilities, P(*Z*_*i*,(*j,k*)_ = 1 | ***Y_i_, π***), which are utilized for assigning each multi-read to the most likely genomic origin. To keep mHi-C output compatible with the input required for the widely used significant interactions detection methods, we filtered multi-reads with maximum allocation posterior probability less than or equal to 0.5 and assigned the remaining multi-reads to their most likely bin pairs. This ensured the use of at most one bin pair for each multi-read pair. We repeated our computational evaluations by varying this threshold on the posterior probabilities to ensure robustness of the overall conclusions to this threshold.

### Assessing false positive rates for significant interactions and TADs identification under the Uni- and Uni&Multi-settings

To quantify false positive rates of the Uni- and Uni&Multi-settings, at the significant interaction level, we defined true positives and true negatives by leveraging deeply sequenced replicates of the IMR90 dataset (replicates 1-4). Significant interactions reproducibly identified across all four replicates at 0.1% FDR by both the Uni- and Uni&Multi-settings were labeled as true positives (i.e., true interactions). True negatives were defined as all the contacts that were not deemed significant at 25% FDR in any of the four replicates. We then evaluated significant interactions identified by smaller depth replicates 5 & 6 with ROC and PR curves (Figure 2–Figure supplement 13) by using these sets of true positives and negatives as the gold standard. To quantify false positive rates at the topologically associating domains (TADs) level (Figure 5C, Figure 5–Figure supplement2E), we utilized TADs that are reproducible in more than three replicates of the IMR90 dataset and/or harbor CTCF peaks at the boundaries as true positives. The rest of the TADs are supported neither by multiple replicates nor by CTCF, hence are regarded as false positives.

### Evaluating reproducibility

Reproducibility in contact matrices was evaluated using HiCRep in the default settings. We further assessed the reproducibility in terms of identified contacts by grouping the significant interactions into three categories: contacts only detected under Uni-setting, contacts unique to Uni&Multi-setting, and contacts that are detected under both settings. The reproducibility is calculated by overlapping significant interactions between every two replicates and recording the percentage of interactions that are also deemed significant in another replicate (Figure 4–Figure supplement 9). Chromatin states of novel significant interactions

### Chromatin states of novel signi!cant interactions

We annotated the novel significant interactions with the 15 states ChromHMM segmentations for IMR90 epigenome (ID E017) from the Roadmap Epigenomics project (***Roadmap Epigenomics Consortium,2015***). All six replicates of IMR90 are merged together in calculating the average enrichment of significant interactions among the 15 states (Figure 4B and Figure 4–Figure supplement 14A).

### ChIP-seq analysis

ChIP-seq peak sets for IMR90 cells were obtained from ENCODE portal (https://www.encodeproject.org/) and GEO (***Barrett et al., 2012***). Specifically, we utilized H3K4me1 (ENCSR831JSP), H3K4me3 (ENCSR087PFU), H3K36me3 (ENCSR437ORF), H3K27ac (ENCSR002YRE), H3K27me3 (ENCSR431UUY) and CTCF (ENCSR000EFI) from the ENCODE project and p65 (GSM1055810), p300 (GSM1055812) and PolII (GSM1055822) from GEO (***Barrett et al., 2012***). In addition, raw data files in fastq format were processed by Permseq (***Zeng et al., 2015***) utilizing DNase-seq of IMR90 (ENCODE accession ENCSR477RTP) to incorporate multi-reads and, subsequently, peaks were identified using ENCODE uniform ChIP-seq data processing pipeline (https://www.encodeproject.org/pages/pipelines/#DNA-binding). CTCF motif quantification for topologically associating domains was carried out with FIMO (***Grant et al., 2011***) under the default settings using CTCF motif frequency matrix from JASPAR (***Khan et al., 2017***).

### Promoters with significant interactions

Significant interactions across six replicates of the IMR90 study were annotated with GENCODE V19 (***Harrow et al., 2012***) gene annotations and enhancer regions from ChromHMM. Gene expression calculations utilized RNA-seq quantification results from the ENCODE project with accession number ENCSR424FAZ.

### Marginal Hi-C tracks in Figure 4–Figure supplements 15-17

Uni-setting and Uni&Multi-setting Hi-C tracks displayed on the UCSC genome browser figures (Figure 4–Figure supplements 15-17) are obtained by aggregating contact counts of six replicates of IMR90 for each genomic coordinate along the genome.

### Visualization of contact matrices and interactions

We utilized Juicebox (***Durand et al., 2016***), HiGlass (***Kerpedjiev et al., 2017***) and WashU epigenome browser (***Zhou et al., 2011***) for depicting contact matrices and interactions, respectively, throughout the paper. Normalization of the contact matrices for visualization was carried out by the Knight-Ruiz Matrix Balancing Normalization (***Knight and Ruiz, 2013***) provided by Juicebox (***Durand et al., 2016***).

### Model-free multi-reads allocation strategies

The simplified and intuitive strategies depicted in Figure 6A correspond to rescuing multi-reads at different essential stages of the Hi-C analysis pipeline. AlignerSelect relies on the base aligner, e.g., BWA, to determine the primary alignment of each individual end of a multi-read pair. DistanceSelect enables the distance prior to dominate. It selects the read end alignments closest in the genomic distance as the origins of the multi-read pair and defaults to the primary alignment selected by base aligner for inter-chromosomal multi-reads. Finally, SimpleSelect follows the overall mHi-C pipeline closely by making use of the standard Hi-C validation checking and binning procedures. For the reads that align to multiple bins, it selects the bin pair closest in the genomic distance as the allocation of the multi-read pair. Bin-pair allocations for inter-chromosomal multi-reads are set randomly in this strategy.

### Trimming procedures

We considered two approaches for generating evaluation datasets where we combined the trimmed multi-reads from replicate 2, which has the median sequencing depth among all replicates of the A549 study set, with (i) trimmed reads of replicate 2 that remain uniquely aligned to the reference genome at the same trimmed read length (Figure 6 and 7), and (ii) uni-reads from other replicates, i.e., replicates 1, 3, and 4 in the A549 dataset individually (Figure 6–Figure supplements 3, 5 and 6). The first setting enables a direct comparison of the set of uni- and multi-reads at trimmed read length compared to uni-reads at the full read length to evaluate accuracy. The numbers of reads are summarized in Figure 6–Figure supplement 1A along with multi-to-uni ratios in Figure 6–Figure supplement 1B. In the second trimming setting (ii), the uni-read sets are of the original sequencing depth and the added multi-reads constitute a smaller proportion compared to observed levels in the data (Figure 6–Figure supplement 1C) due to the chimeric read rescue that was part of full-length datasets (Figure 7–Figure supplement 8). Therefore, for this setting, we leverage the higher overall depth of the datasets and evaluate the multi-read assignment accuracy at different resolutions, i.e., 10kb and 40kb.

### Simulation procedures

We devised a simulation strategy that utilizes parameters learned from the Hi-C data and results in data with a similar signal to noise characteristics as the actual data.

1. *Construction of the interaction prior based on the uni-reads fragment interaction frequency list of GM12878 dataset (replicate 6)*. The frequency list from the prior encompasses both the genomic distance effect and local interaction signal strength and forms the basis for simulating restriction fragment interactions.
2. *Generating the restriction enzyme cutting sites for each simulated fragment pair*. After sampling interacting fragments using the frequency list from Step 1, a genomic coordinate within ± 500bp of the restriction enzyme cutting site and a strand direction are selected randomly. Reads of different lengths (36bp, 50bp, 75bp, 100bp) are generated starting from these cutting sites.
3. *Mutating the resulting reads*. Mutation and gap rates are empirically estimated based on the aligned uni-reads of replicate 6. The reads from Step 2 are uniformly mutated with these rates allowing up to 2 mutations and 1 gap.
4. *Simulate sequence quality scores of the reads*. We utilize the empirical estimation of the distribution regarding the sequence base quality scores across individual locations of the read length and simulate for each read its sequence quality scores at the nucleotide level.
5. *Alignment to the reference genome*. The simulated reads are aligned to the reference genome and filtered for validation as we outline in the mHi-C pipeline, resulting in the set of multi-reads that are utilized by mHi-C.

We generated numbers of multi-reads comparable to those of replicate 6 (Figure 7–Figure supplement 4B) and in the final step of the simulation studies, we merge the simulated set of multi-reads with rep3&6 uni-reads and ran mHi-C step4 (binning) - step5 (prior already available) - step6 (assign multi-reads posterior probability) independently at resolutions 10kb and 40kb.

### Software availability

mHi-C pipeline is implemented in Python and accelerated by C. The source codes and instructions for running mHi-C are publicly available at https://github.com/keleslab/mHiC. Each step is organized into an independent script with flexible user-defined parameters and implementation options. Therefore, analysis can be carried out from any step of the work-flow and easily fits in high-performance computing environments for parallel computations.

## Acknowledgments

This work was supported by NIH HG009744 and NIH HG007019 (S.K.). F.A. is partially supported by Institute Leadership Funds from La Jolla Institute for Allergy and Immunology. We thank Peigen Zhou from the University of Wisconsin-Madison for the insightful discussions on accelerating the pipeline. We also thank the peer reviewers and the Reviewing and Senior ***eLife*** Editors of this work for their constructive comments.

## Author Contributions

F.A. and S.K. conceived the project. Y.Z., F.A., and S.K. designed the research. Y.Z. and S.K. developed the method. Y.Z. constructed the pipeline and performed experiments. All authors contributed to the preparation of the manuscript.

---

**Figure 1–source data 1.** Detailed summary of study datasets.

**Figure 1–Figure supplement 1.** mHi-C pipeline (Alignment - Read end pairing - Valid fragment filtering).

**Figure 1–Figure supplement 2.** mHi-C pipeline (Duplicate removal - Genome binning - mHi-C).

**Figure 1–Figure supplement 3.** Coverage and cis-to-trans ratios across individual replicates of the study datasets as indicators of data quality.

**Figure 1–Figure supplement 4.** Percentages of mappable and valid reads across study datasets as an indicator of data quality.

**Figure 1–Figure supplement 5.** Categorization of reads after alignment across study datasets.

**Figure 1–Figure supplement 6.** Comparison of the prevalence of multi-reads and chimeric reads, both of which require additional processing.

**Figure 2–Figure supplement 1.** Raw and normalized contact matrices of GM12878 under Uni-setting and Uni&Multi-setting fon chromosome 1.

**Figure 2–Figure supplement 2.** Raw and normalized contact matrices of GM12878 under Uni-setting and Uni&Multi-setting on chromosome 2.

**Figure 2–Figure supplement 3.** Raw and normalized contact matrices of GM12878 under Uni-setting and Uni&Multi-setting on chromosome 3.

**Figure 2–Figure supplement 4.** Raw and normalized contact matrices of GM12878 under Uni-setting and Uni&Multi-setting on chromosome 5.

**Figure 2–Figure supplement 5.** Bin coverage improvement of Uni&Multi-setting compared to Uni-setting for IMR90 at the individual replicate level for two different allocation probability thresholds.

**Figure 2–Figure supplement 6.** Reproducibility at the contact matrix level under the Uni- and Uni&Multi-settings across study datasets.

**Figure 2–Figure supplement 7.** Reproducibility at the contact matrix level at resolutions 40kb (low) and 10kb (high) across study datasets.

**Figure 2–Figure supplement 8.** Improvement in reproducibility versus multi-read contribution across chromosomes.

**Figure 2–Figure supplement 9.** A detailed analysis of reproducibility of significant interactions for IMR90.

**Figure 3–source data 1.** Percentage of improvement in the number of significant interactions across 6 studies at resolution 40kb.

**Figure 3–source data 2.** ChIP-seq peaks detected following the standard ChIP-seq data processing pipeline of ENCODE (***The ENCODE Project Consortium, 2012***) using both uni-reads and multi-reads aligned by Permseq (***Zeng et al., 2015***) for CTCF, PolII, p65, p300, H3K27ac, H3K27me, H3K36me3, H3K4me1 and H3K4me3.

**Figure 3–Figure supplement 1.** Percentage change in the numbers of significant interactions under the Uni&Multi-setting compared to Uni-setting at different FDR thresholds and resolutions.

**Figure 3–Figure supplement 2.** Comparison of significant interactions as a function of posterior probabilities of multi-read assignment (IMR90).

**Figure 3–Figure supplement 3.** Heatmap for marginal correlations of percentage increase in the number of identified significant interactions (FDR 5%) with indicators of data quality.

**Figure 3–Figure supplement 4.** Percentage change in the numbers of significant interactions with respect to cis-to-trans ratio.

**Figure 3–Figure supplement 5.** Percentage change in the numbers of significant interactions of GM12878 datasets at different resolutions.

**Figure 3–Figure supplement 6.** Percentage change in the numbers of significant interactions with respect to coverage.

**Figure 3–Figure supplement 7.** Percentage change in the numbers of significant interactions as a function of the percentage of mHi-C rescued multi-reads in comparison to uni-reads and cis-to-trans ratio.

**Figure 3–Figure supplement 8.** Recovery of significant interactions identified at FDR 1% by analysis at FDR 10% for each of six replicates of IMR90.

**Figure 3–Figure supplement 9.** Recovery of significant interactions identified at FDR 1% by analysis at FDR 10% for each of four replicates of GM12878 at 40kb resolution.

**Figure 3–Figure supplement 10.** Recovery of significant interactions identified at FDR 1% by analysis at FDR 10% for each of four replicates of GM12878 at 10kb resolution.

**Figure 3–Figure supplement 11.** Recovery of significant interactions identified at FDR 1% by analysis at FDR 10% for each of ten replicates of GM12878 at 5kb resolution.

**Figure 3–Figure supplement 12.** Recovery of significant interactions identified at FDR 1% by analysis at FDR 10% for GM12878 aggregated across replicates at 5kb, 10kb, and 40kb resolutions.

**Figure 3–Figure supplement 13.** ROC and PR curves for replicates 5 and 6 of IMR90.

**Figure 3–Figure supplement 14.** Quantification of significant interactions for chromHMM states and ChIP-seq peak regions (IMR90).

**Figure 3–Figure supplement 15.** Marginalized Hi-C signal, ChIP-seq coverage and peaks, and gene expression for chr1 (IMR90).

**Figure 3–Figure supplement 16.** Marginalized Hi-C signal, ChIP-seq coverage and peaks, and gene expression for chr2 (IMR90).

**Figure 3–Figure supplement 17.** Marginalized Hi-C signal, ChIP-seq coverage and peaks, and gene expression for chr9 (IMR90).

**Figure 4–source data 1.** The number of significant promoter-enhancer Hi-C interactions at FDR 5% under Uni-setting and Uni&Multi-setting, respectively, for six replicates of IMR90.

**Figure 4–source data 2.** Significant promoter-enhancer interactions at FDR 5% under Uni-setting and Uni&Multi-setting for six replicates of IMR90 with the number of contacts.

**Figure 4–Figure supplement 1.** Examples of significant promoter-enhancer interactions reproducible among 6 replicates under Uni- and Uni&Multi-settings (IMR90) on chromosome 7.

**Figure 4–Figure supplement 2.** Examples of significant promoter-enhancer interactions reproducible among 6 replicates under Uni- and Uni&Multi-settings (IMR90) on chromosome 17.

**Figure 4–Figure supplement 3.** Significant promoter-enhancer interactions under Uni- and Uni&Multi-settings across 6 IMR90 replicates (Chromosome 17).

**Figure 4–Figure supplement 4.** Expression distribution of genes promoters of which have significant promoter interactions (IMR90).

**Figure 5–source data 1.** Topologically associating domains detected by DomainCaller (***Dixon et al., 2012***) under Uni&Multi-setting for six replicates of IMR90.

**Figure 5–source data 2.** Topologically associating domains detected by Arrowhead (***Rao et al., 2014***) under Uni&Multi-setting for ten replicates of GM12878.

**Figure 5–Figure supplement 1.** The number of topologically associating domains (TADs) detected in each chromosome under Uni-setting and Uni&Multi-setting (IMR90).

**Figure 5–Figure supplement 2.** Comparison of CTCF peaks at the boundaries of topologically associating domains (TADs) under Uni-setting and Uni&Multi-setting across six replicates of IMR90.

**Figure 5–Figure supplement 3.** Novel topologically associating domains (TADs) with CTCF peaks at TAD boundaries (IMR90).

**Figure 5–Figure supplement 4.** Existing topologically associating domains (TADs) with adjusted boundaries supported by CTCF peaks at the new TAD boundaries on chr1 and chr5 (IMR90).

**Figure 5–Figure supplement 5.** Existing topologically associating domains (TADs) with adjusted boundaries supported by CTCF peaks at the new TAD boundaries on chr12 and chr13 (IMR90).

**Figure 5–Figure supplement 6.** False positive topologically associating domains (TADs) detected by the Uni-setting due to the missing reads in low mappability regions on chr2 and chr3 (IMR90).

**Figure 5–Figure supplement 7.** False positive topologically associating domains (TADs) detected by the Uni-setting due to the missing reads in low mappability regions on chr4 and chr16 (IMR90).

**Figure 5–Figure supplement 8.** False positive topologically associating domains (TADs) detected by the Uni-setting due to the missing reads in low mappability regions on chr21 and chrX (IMR90).

**Figure 5–Figure supplement 9.** False discovery rate of TADs detected under two settings. TADs that are not reproducible are labeled as false positives without considering the CTCF peaks at the TAD boundaries.

**Figure 5–Figure supplement 10.** Percentage of TAD boundaries harboring different types of repetitive elements under Uni-setting and Uni&Multi-setting for IMR90 at 40kb and GM12878 at 5kb.

**Figure 5–Figure supplement 11.** Average number of repetitive elements at the reproducible TAD boundaries compared to averages within TADs and genomewide intervals of the same size for IMR90 at 40kb resolution.

**Figure 6–Figure supplement 1.** Summary of the sequencing depths of the full length and trimmed datasets of the A549 study.

**Figure 6–Figure supplement 2.** Allocation accuracy at the 40kb resolution among different mappability regions for trimmed reads of varying lengths.

**Figure 6–Figure supplement 3.** Intra-chromosomal and intra&inter-chromosomal allocation accuracy with respect to trimmed read length for trimming setting (ii).

**Figure 6–Figure supplement 4.** Evaluating accuracy of mHi-C multi-read allocation with simulations.

**Figure 6–Figure supplement 5.** Allocation accuracy across different mappability regions for trimmed reads of length 36bp, 50bp, 75bp, 100bp, and 125bp for trimming setting (ii).

**Figure 6–Figure supplement 6.** Allocation accuracy across different classes of repetitive elements at 10kb and 40kb resolutions under trimming setting (ii).

**Figure 6–Figure supplement 7.** Comparison of significant interactions among the three life stages of *P. falciparum*.

**Figure 7–Figure supplement 1.** Reproducibility of trimmed Uni-setting and trimmed Uni&Multi-setting across different read lengths at 40kb resolution.

**Figure 7–Figure supplement 2.** TAD detection on chromosome 1 of original replicate uni-reads contact matrix, trimmed uni-reads (36bp) contact matrix and trimmed uni- and multi-reads (36bp) contact matrix.

**Figure 7–Figure supplement 3.** TAD detection on chromosome 4 of original replicate uni-reads contact matrix, trimmed uni-reads (50bp) contact matrix and trimmed uni- and multi-reads (50bp) contact matrix.

**Figure 7–Figure supplement 4.** TAD detection on chromosome 7 of original replicate uni-reads contact matrix, trimmed uni-reads (75bp) contact matrix and trimmed uni- and multi-reads (75bp) contact matrix.

**Figure 7–Figure supplement 5.** TAD detection on chromosome 10 of original replicate uni-reads contact matrix, trimmed uni-reads (100bp) contact matrix and trimmed uni- and multi-reads (100bp) contact matrix.

**Figure 7–Figure supplement 6.** Numbers of significant interactions identified with trimmed reads under Uni- and Uni&Multi-settings at FDR 0.1%, 1%, 5%, 10%.

**Figure 7–Figure supplement 7.** Proportion of top 10,000 significant interactions of full read length dataset recovered in trimmed reads sets under FDR 10%.

**Figure 7–Figure supplement 8.** Impact of chimeric reads in the A549 datasets.

**Figure 7–Figure supplement 9.** ROC and PR curves for detection of full read length dataset significant interactions by the analysis of trimmed read datasets.

